# Production and secretion of recombinant proteins using the endotoxin-free, Gram-negative bacterium *Sphingobium japonicum*

**DOI:** 10.1101/2022.09.04.506512

**Authors:** Ehud Shahar, Ken Emquies, Jacob Pitcovski, Itai Bloch, Dalia Eliahu, Ran Ben Adiva, Itamar Yadid

**Affiliations:** Migal – Galilee Research Institute Kiryat Shmona - Israel; Tel-Hai Academic College Kiryat Shmona – Israel

**Keywords:** *Sphingomonas*, Recombinant subunit vaccine, Lipopolysaccharide-free, Glycosphingolipids, Signal peptides

## Abstract

Gram-negative bacteria are common and efficient protein expression systems, yet their outer membrane endotoxins can elicit undesirable toxic effects, limiting their applicability for parenteral therapeutic applications, e.g., production of vaccine components. In the bacterial genus *Sphingomonas* from the *Alphaproteobacteria* class, lipopolysaccharide (LPS) endotoxins are replaced with non-toxic glycosphingolipids (GSL), rendering it an attractive alternative for therapeutic protein production. To explore the use of *Sphingomonas* as a safe expression system for production of proteins for therapeutic applications, in this study, *Sphingobium japonicum* (*SJ*) injected live into embryonated hen eggs proved safe and nontoxic. Multimeric viral polypeptides derived from Newcastle disease virus (NDV) designed for expression in SJ, yielded soluble proteins which were specifically recognized by antibodies raised against the whole virus. In addition, native signal peptides (SP) motifs identified using whole-genome computerized analysis and coupled to secreted proteins in *SJ* induced secretion of αAmy and mCherry gene products. Relative to the same genes expressed without an SP, SP 104 increased secretion of αAmy (3.7-fold) and mCherry (16.3-fold) proteins and yielded accumulation of up to 80µg/L of the later in the culture medium. Taken together, the presented findings demonstrate the potential of this unique LPS-free Gram-negative bacterial family to serve as an important tool for protein expression for both research and biotechnological purposes, including for the development of novel vaccines.

**Importance:** The Gram-negative, LPS-free *Sphingomonas* genus can serve as an effective and safe platform for therapeutic protein expression and secretion and bears potential as a live bacteria delivery system for protein vaccines.

## Background

*E. coli* is the main bacterial species used for protein expression for research, industrial and pharmaceutical applications. The comprehensive data available regarding its genetic and physiological characteristics offer an abundance of tools together with well-established work protocols. These call for use of readily available, simple and inexpensive media, and enable genetic manipulation, rapid propagation and high target protein yields (Schmidt 2004; Rosano and Ceccarelli 2014; Zhang et al. 2018). Despite the said advantages, protein expression in *E. coli* can generate misfolded proteins that lack both activity and conformational epitopes required to produce neutralizing antibodies in vaccines. Improper folding in a prokaryote-based system can also result in the accumulation of inclusion bodies (IB) or no expression at all. Moreover, sample contamination with the highly toxic *E. coli* lipopolysaccharide (LPS) requires a critical purification step (Petsch and Anspach 2000; Kawahara et al. 1999; Alexander and Rietschel 2004).

The *Sphingomonadaceae* family is a unique Gram-negative bacteria, best known for its ability to degrade a wide variety of man-made pollutants including dioxin, biphenyl, bisphenol, and pentachlorophenol (Ederer et al. 1997; Prakash and Lal 2006), and to produce complex biopolymers, namely gellan gum and additional related polysaccharides (Krziwon et al. 1995). While molecular tools for genetic manipulation of and protein overexpression in this family are still limited, a growing number of studies is being conducted to further establish and expand *Sphingomonadaceae* applications in biotechnology (Heaver et al. 2018; Lin et al. 2021).

In contrast to other Gram negative bacteria, members of the *Sphingomonadaceae* family do not generate LPS, but rather, produce the much less immunogenic molecule - glycosphingolipids (GSL), which structurally and physicochemically resembles the lipid A component of LPS (Kawahara et al. 1999). The biological role of GSL is still not fully understood and is being actively researched (Olsen and Jantzen 2001). Studies have shown that some GSLs induce immunological activity 10,000-fold lower compared to LPS whereas others are entirely inactive, with no immunotoxic effects (Krziwon et al. 1995). In addition, *Sphingomonadaceae* bacteria do not regularly colonize or invade tissues and are rarely associated with infectious diseases despite their widespread existence in the environment (Arora and Porcelli 2008). Furthermore, as a result of LPS replacement by GSL, which form unique membrane properties, their ability to fold and secrete challenging proteins might be improved. Considering these characteristics, *Sphingomonadaceae* may offer an attractive bacterial platform for protein expression and production.

The current study evaluated the potential of *Sphingobium japonicum* (SJ), a member of the *Sphingomonas* family, to serve as a live protein delivery vector, and to produce and secrete polypeptides aimed for therapeutics and vaccination.

## Materials and Methods

### Toxicity evaluation following *in-ovo* injection of bacteria

*E. coli* strain BL21 (DE3) and *S. japonicum* UT26 (NBRC 101211) were grown to OD600=1, which is equivalent to 5×10^8^ cells/ml. *E. coli, S. japonicum* or PBS as control, were injected (100μl) into the allantoic fluid of embryonated specific pathogen-free eggs (Charles River, Wilmington, MA) preincubated for 18 days (37 °C, ∼80% relative humidity and gentle rocking) (Fig. 1). Eggs were further incubated, and hatching percentage was determined on day 21, i.e., 3 days after injection). The hatched chicks were monitored for an additional 14 days and inspected for signs of morbidity.

**Figure 1.**
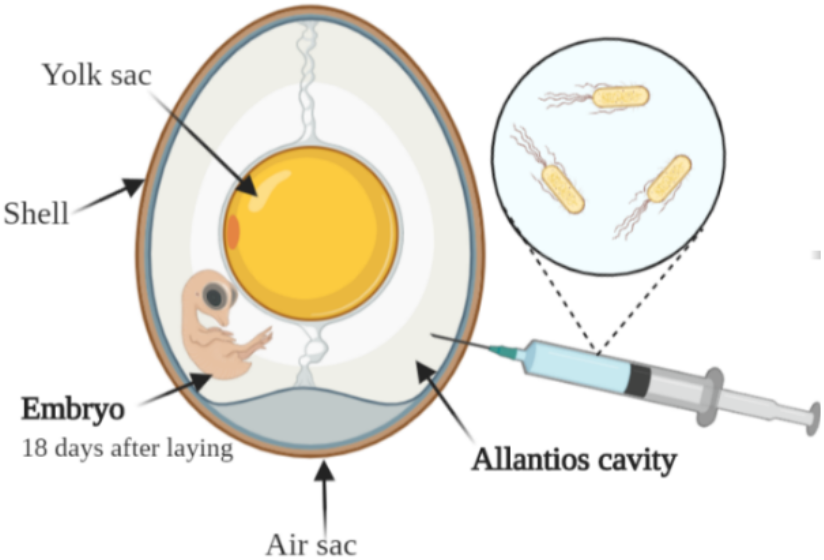
Effect of *in-ovo* bacterial injection on embryonated hen egg hatching rate. To assess the potential immunological advantage of SJ to serve as a vector and expression system, live bacteria were injected into hen eggs. The bacterial cells were suspended in PBS and 0.1 ml of cultures of similar optical densities (OD_600_=1) were injected into the allantois cavity on Day 18 of incubation (n is indicated above each treatment bar). PBS: control group, *E. coli*: BL21 (DE3) strain, *S. japonicum*: SJ wild type. Illustration was drawn using BioRender.

### Native signal peptide selection

The genome sequence of *S. japonicum* UT26 was from NCBI GenBank AP010803-6 (Nagata et al. 2010). All open reading frames (ORFs) in the genome were defined via NCBI ORFinder and translated into amino acid sequences. All sequences were submitted to the SignalP 5.0 server for prediction of Sec secretory signal peptides (SPs) (Emanuelsson et al. 2007). Out of the 200 top-scoring SP sequences, six representatives (SP1, SP2, SP5, SP99, SP104 and SP199) were selected for further analysis (Table 1).

**Table 1.**
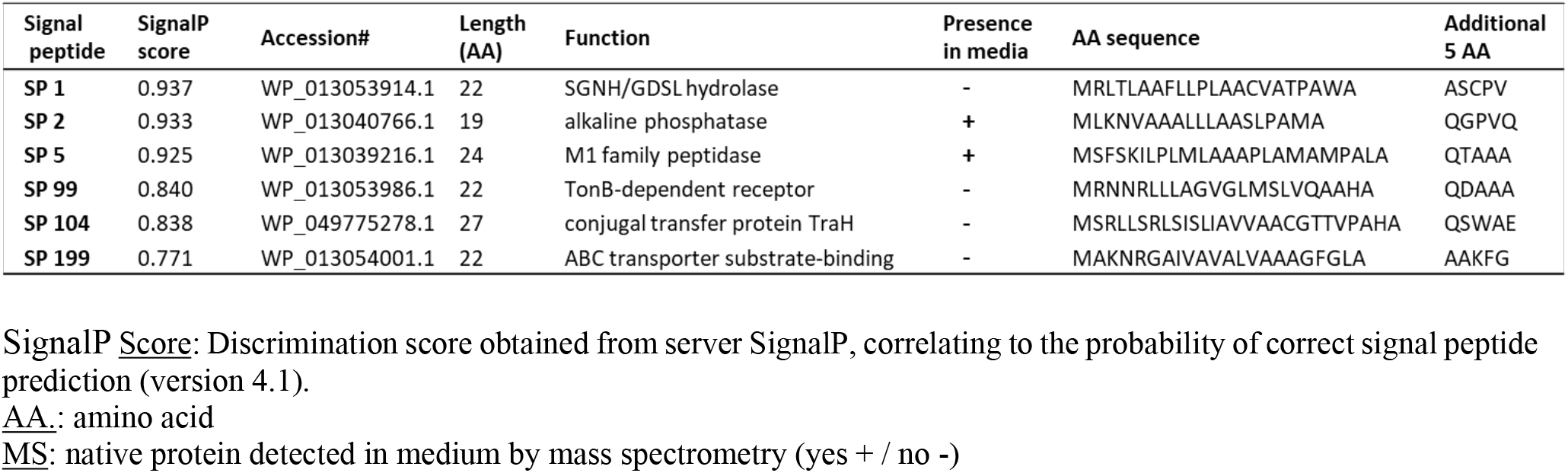
The predicted native signal peptides obtained from server SignalP.

### Identification of proteins naturally secreted to the medium by *S. japonicum*

To identify the proteins secreted into the medium, cells were pelleted, the supernatant was collected and filtered and then subjected to TCA precipitation, as previously described (Sanchez 2001). The precipitate was dried by incubation at 95 °C for 10 min, resuspended in 100 μl 1X sample buffer containing beta-mercaptoethanol and then heated for 5 min at 95 °C. Samples were analyzed by mass spectrometry (MS) at the Smoler Proteomics Center at the Technion Institute, Haifa, Israel.

### Plasmid construction

The α amylase (αAmy) gene was amplified from the genomic DNA of *B. licheniformis* (NBRC 12200) using the following primers: forward 5’-CTAGTAGAGGAAGCTTCCGCATGCTCGAGGCAAATCTTAAAGGGACGCTG-3’, reverse-5’-GTAGTCCGGATCCCAATTGGAGCTCCTATCTTTGAACATAAATTGAAACCGAC-3’. The αAmy gene was cloned into pVH plasmid vector (Kaczmarczyk et al. 2014)using a restriction-free (RF) cloning technique, according to a previously published protocol (Van Den Ent and Löwe 2006). The mCherry gene in the plasmid pVCC was obtained from Addgene (Kaczmarczyk et al. 2014), and digested with XhoI and EcoRI and cloned into pVH digested with the same enzymes. All SP sequences were synthesized by Genscript® and inserted into the expression vectors via restriction with HindIII and XhoI (Thermo Scientific™ FastDigest) followed by ligation (Fig. 2). To preserve the proteolytic cleavage site, the native 5 amino acids (AA) downstream to the SP sequence were included (Auclair et al. 2012). *E. coli* DH5α (ATCC 53868) was used for all plasmid amplifications.

**Figure 2.**
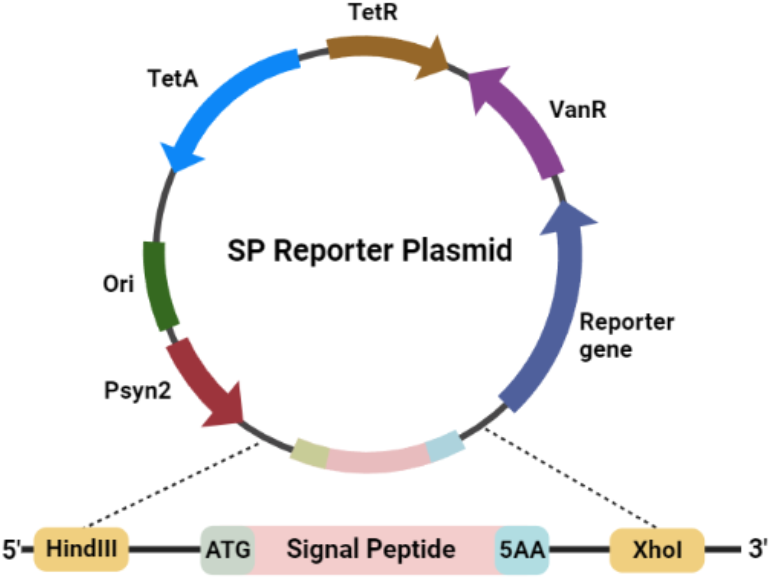
Signal peptide reporter plasmid map and signal peptide integration. The signal peptide (SP) sequence was designed for insertion into a pVH expression vector. Restriction sites HindIII and XhoI were inserted at the 5’ and 3’ ends, respectively. ATG: start codon, 5AA: five amino acids downstream to the SP of the original gene.

### *SJ* transformation

SJ colonies were grown in 5 ml four-fold diluted tryptic soy broth (1/4 TSB) medium overnight (30 °C, 250 rpm) and then transferred to 45 ml fresh 1/4 TSB and grown until O.D600nm=0.6. The cells were pelleted, rinsed with water, and then suspended in 1 ml 300 mM sucrose solution (dissolved in double distilled water (DDW)). Then, 0.25-1 µg plasmid was added to 100 µl competent cells and incubated on ice for 5 min. The mixture was transferred to a pre-chilled 2 mm gap electroporator cuvette and cells were electroporated using the EC2 program (MicroPulser, Bio-Rad). Cells were then suspended in 3 ml 1/4 TSB (at room temperature, rt) supplemented with 0.2 % glucose and grown (30 °C, 250 rpm) overnight. Cells (100 µl) were then seeded on selective plates containing 1/4 TSB agar supplemented with 12.5 µg/ml tetracycline, 10 µg/ml piperacillin, and 0.2% glucose and further incubated for 2-3 days, at 30 °C. Plasmid presence in forming colonies was verified by polymerase chain reaction (PCR).

### Protein expression in SJ

Individual PCR-verified colonies were incubated overnight (30 °C, 250 rpm) in 10 ml selective 1/4 TSB medium supplemented with 12.5 µg/ml tetracycline, 10 µg/ml piperacillin, and 0.2% glucose. Cultures were diluted 25-fold into fresh selective medium and incubated until O.D600 reached 0.8-1. Protein expression was then induced by incubating cells for 3-4 h, at 30 °C, in the presence of 250 μM vanillate.

### Alpha-amylase activity assays

To test the ability of the predicted SPs to induce the secretion of αAmy on solid medium, α amylase activity was evaluated in a Petri dish. Three colonies from each clones bearing a SP-αAmy construct were grown in 1 ml of selective medium until O.D600 reached 0.6. Then, 10 µl of each culture were spotted in triplicates on selective agar plates supplemented with 0.2 % soluble starch and 250 µM vanillate and incubated overnight at 30 °C. Starch hydrolysis was visualized and evaluated after staining the plate with KI/I2 solution for 10 min at room temperature.

αAmy secreted into liquid medium was assayed by gel zymography. Growth medium (10 ml) from each clone carrying a SP-αAmy construct was collected and concentrated four-fold. The samples were mixed with nonreducing sample buffer and loaded onto 10% SDS PAGE gel containing 0.2% soluble starch. Purified *B. licheniformis* αAmy (1 mg/ml) was loaded as a positive control. The gel was then placed in a renaturing buffer (1% Triton x-100) for 10 min, with gentle agitation, and rinsed five times in DDW. The gel was then incubated in activity buffer (50 mM Tris-HCl pH 7 and 1 mM CaCl2) for 3 h, at room temperature, on an orbital shaker. Finally, the gel was stained with KI/I2 solution until the appearance of clear bands.

To quantify the ability of the various SPs to induced secretion of αAmy protein in liquid medium, four colonies for each SP were grown overnight in 2 ml selective medium and then centrifuged at 3,000 rpm, 4°C for 10 min. The cells were resuspended in 10 ml LB supplemented with 12.5 µg/ml tetracycline, 10 µg/ml piperacillin and incubated (30 °C, 250 rpm) until O.D600 reached 0.6. Thereafter, vanillate was added (to 250 µM) and cells were further incubated for 4 h, and then centrifuged at 3,500 rpm for 15 min, at 4°C. To measure α-amylase activity 50 µl of the supernatant was mixed with 50 µl starch solution (20 mM sodium phosphate pH 6.9, 6.7 mM NaCl, 0.2 % soluble starch, and 1 mM CaCl2) and incubated for 2 h, at 55 °C. Then, samples were mixed with 100 µl 3,5-dinitrosalicyclic acid (DNS) and incubated at 95 °C for 5 min before the absorbance was measured at 540 nm using a microplate spectrophotometer (BioTek HT).

### Quantification of mCherry secretion

SJ colonies carrying an expression plasmid with the various SPs followed by the mCherry gene were grown in 10 ml selection medium to an O.D600 of 0.8. Expression was induced as described above for αAmy. Then, the cells were centrifuged at 3500 rpm for 15 min at 4 °C and the medium was collected and concentrated four-fold. Protein concentration in the growth medium was quantified by measuring fluorescence at 610 nm following excitation at 575 nm in comparison of signals with a calibration curve prepared from mCherry which was cloned into pQE30 (NEB), expressed in *E. coli* DH5α and purified using zinc affinity chromatography.

### Cloning of Newcastle disease virus (NDV)-derived polypeptides

The NDV envelop proteins hemagglutinin-neuraminidase Uniprot-P35743 (HN) and the fusion protein Uniprot-P33614 (F) form the LaSota strain were used to design the expression constructs together with the available protein structures (PDB 1G5G and 4FZH respectively (Chen et al. 2001; Yuan et al. 2012)). The genes for the designed HN protein variant 1 (HNv1) and F protein variant 1 (Fv1) were codon-optimized for expression in SJ and synthesized by Genscript®. The genes with or without SP 104 were cloned into the pVH plasmid as described above, using HindIII and XhoI restriction enzymes.

### Detection of expressed NDV polypeptides by western blot (WB) analysis

Following the induction of NDV polypeptide expression in SJ, cells were separated from the growth medium by centrifugation, then lysed by sonication and centrifuged at 10,000 g at 4 °C. Lysate pellet and lysate supernatant were loaded onto SDS-PAGE gels and assayed by western blotting as follows. SDS-PAGE gels were washed twice in 30 ml DDW for 5 min, and proteins were then transferred to a nitrocellulose membrane using a Trans-Blot Turbo Nitrocellulose Transfer Pack and Transfer System (Bio-Rad). The membrane was blocked for 1 h with 5% skim milk in PBS supplemented with 0.05% Tween 20 (PBS-T) and then incubated for 1 h with rabbit anti polyhistidine-HRP or chicken serums (1:200), followed by 1 h incubation with goat anti-chicken IgY-HRP (Thermo Scientific). The membrane was washed three times for 5 min with PBS-T buffer after each incubation. Finally, the membrane was placed in 500 µl of Clarity Western ECL Blotting Substrate and Enhancer (Bio-Rad) and chemiluminescence was detected using an ImageQuant™ LAS 4000.

### Chicken vaccination with HNv1 expressed in SJ

Cell pellets from 50 ml cultures of induced HNv1-expressing SJ cells were resuspended in 5 ml PBS and lysed by sonication. Cell lysates (1.75 ml) were then diluted 2-fold in PBS and mixed with 3.5 ml complete Freund’s adjuvant or incomplete Freund’s adjuvant for the first and second vaccine doses, respectively (Thermo Scientific). Adjuvanted PBS was prepared as a negative control, and commercial inactivated viral vaccine (VH, Phibro™) was used as a positive control. Chickens were maintained on a 24 °C, 12:12 light:dark regimen and food ad lib. Two vaccine doses (1 ml) were injected half subcutaneously and half intramuscularly, with the first dose administered at 3 weeks of age and the second 2 weeks later. Blood was drawn from the jugular vein 2 weeks after the last injection. Sera were extracted and stored at -20 °C until use.

### Enzyme-linked immunosorbent assay (ELISA)

Sandwich ELISA was performed with mouse anti-NDV-HN antibodies (OriGene) diluted 1:100 in PBS for 1 h to capture whole attenuated NDV (VH, Phibro™). The plate was blocked (1 h, room temperature) with 300 μl PBS-T supplemented with 5% skim milk. Following NDV capture for 1h at RT and PBS wash, serum was added and then detected with goat anti-chicken IgY-HRP (Thermo Scientific). Plates were then incubated with TMB substrate solution (Thermo Scientific), and reaction was terminated by addition of sulfuric acid. The absorbance was measured at 450 nm using a microplate spectrophotometer (BioTek HT).

### Statistical analysis and illustrations design

Statistical analysis and graph design were performed using GraphPad Prism version 9 (GraphPad Software Inc, USA). ANOVA test with Tukey post hoc analysis was performed to determine significant differences between groups. Illustrations were designed using the BioRender website version.

## Results

### Toxicity of SJ in embryonated eggs

To evaluate the potential toxic effect of LPS-free *SJ*, the allantois cavity of fertilized hen eggs (n=6) was injected with an equal amount of SJ or *E. coli* BL21 cells, which served as an LPS-positive control. The injection was performed on day 18 (out of 21) of egg incubation, when the embryo’s adaptive immune system is functional, and eggs are routinely transferred from an incubator to a hatcher (Alqhtani et al. 2022). Following *in ovo* administration of *E. coli*, 0/6 chicks hatched, whereas all chicks injected with SJ hatched after 21 days and survived for at least two weeks, with no apparent side effects or abnormal signs. Similar hatching rates were noted in the group injected with PBS.

### Identification, design, and cloning of native secretion signal peptides

To identify potential signal peptides (SPs), the genome of SJ was screened for protein sequences containing SP motifs. Out of the first 200 predicted native SPs, six were selected for additional characterization following validation of presence of a putative secreted protein sequence downstream (Table 1).

In addition, MS analysis of proteins secreted following growth of SJ in rich medium identified over 300 proteins. Out of the 6 selected SP sequences, only SP 2 and SP 5 which drives the secretion of alkaline phosphatase and the M1 family protease respectively, were detected in culture medium by the MS analysis (Table 1). To evaluate the ability of the six selected SP sequences to drive the secretion of model proteins, the SP sequences were inserted into a pVH expression vector carrying the α Amy and mCherry reporter genes (Figure 2). All but one SP construct (SP 99) yielded positive colonies. Thus, all further experiments were carried out using the remaining five SPs (Table 1).

### Influence of addition of SP sequence on the secretion of αAmy and mCherry model proteins

The ability of the predicted SPs to drive secretion of active αAmy was examined by growing colonies transformed with a vector carrying different SP sequences upstream to the αAmy gene on plates supplemented with starch. SP 2, SP 5, and SP 104 were associated with strong α amylase activity, evident by the formation of large yellow/orange halos around all colonies tested, while αAmy cloned with SP 1 and SP 199 were associated with a relatively weak or practically undetectable signal, respectively (Figure 3A).

**Figure 3.**
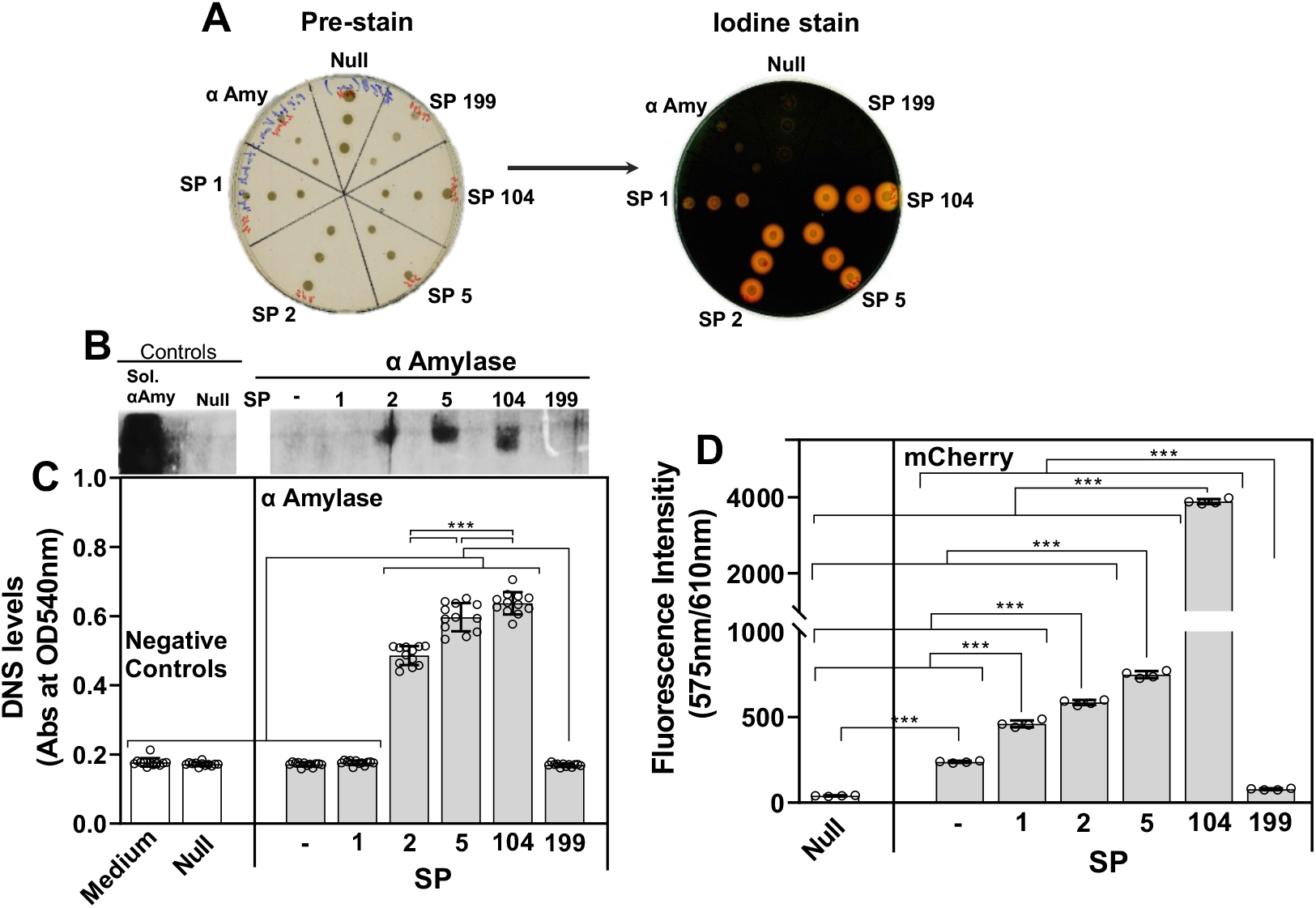
The influence of signal peptides on protein secretion in *Sphingobium japonicum*. **A**. Secretion of α-Amy was tested in solid medium. Three colonies from each SJ transformed with a pVH vector expressing either signal peptide (SP) 1, SP 2, SP 5, SP 104, SP 199, Null (pVH empty vector) or αAmy (pVH vector with α-amylase without addition of SP) were grown on 0.2% starch Petri dishes (left). Iodine stain solution was added to detect starch hydrolysis (right). Secretion of αAmy (B, C) and mCherry (D) into liquid medium was tested and the activity quantified by using growth medium collected from SJ cells carrying empty pVH vector as negative control (Null) and growth medium collected from SJ cells carrying the αAmy gene without (-) or with SP 1, SP 2, SP 5, SP 104, or SP 199. **B**. Negative image visualization of gel zymography of purified αAmy sample as positive control (Sol. αAmy). **C**. Mean ± SD (n=12) α amylase activity as measured by starch degradation into reducing sugars, tested using the 3,5-dinitrosalicyclic acid (DNS) assay. **D**. Mean ± SD (n=4) mCherry fluorescence intensity (575nm/610nm) measured in growth medium. One-way ANOVA with Tukey’s post hoc was applied to determine statistically significant differences (*** p≤0.001).

Quantification of αAmy and mCherry secretion in liquid medium following induction found similar results as observed in solid medium, with high α amylase activity in samples bearing SP 2, SP 5, and SP 104 (Figure 3B). No activity was observed in proteins linked to SP 1, SP 199 or lacking a SP altogether. Further quantification of the secreted α amylase activity using the DNS assay found the highest enzymatic activity in the medium when αAmy secretion was driven by SP 104, with 3.7-fold higher levels compared to samples with no SP (p<0.001). Activity of αAmy cloned with SP 2 and SP 5 was also significantly higher than that measured in colonies lacking SP (2.8-fold and 3.4-fold increase, respectively; Figure 3C).

mCherry fluorescence signals followed a similar trend. SP 104-driven secretion was associated with a 16.3-fold higher fluorescence signal as compared to samples lacking a SP (p<0.001). SP 1, SP 2 and SP 5 also significantly increased the measured fluorescence signal as compared to mCherry secreted without addition of SP. Notably, the addition of SP 199 sequence to the mCherry gene resulted in a 3.2-fold decreased signal as compared to cells carrying the gene without a SP (Figure 3D). Under the tested conditions, SP 104, which was the most potent SP, induced secretion of 80.6μg/L mCherry (Table 2).

**Table 2.** Quantification of mCherry secreted in broth medium.

| Signal peptide | Fluorescence (a.u) | MCherry concentration (μg/ml) |
| --- | --- | --- |
| SP 2 | 585.5 | 12.2 |
| SP 5 | 748 | 15.6 |
| SP 104 | 3885.25 | 80.6 |

a.u.: arbitrary units.

### Design and construction of a subunit vaccine against Newcastle Disease Virus

Proteins sequences derived from the NDV virus, a prominent virus infecting wild and domestic birds were used as a model for heterologous protein production and to evaluate the therapeutic potential of proteins produced in SJ and for vaccination. NDV expresses fusion protein (F) and hemagglutinin neuraminidase protein (HN), two important functional surface glycoproteins which induce the production of neutralizing antibodies. The F protein is a homotrimer, synthesized as a non-fusogenic F0 proprotein and is converted to the fusogenic protein after proteolytic cleavage by a host cell protease. The mature protein consists of two disulfide-linked subunits: F1 and F2. The F protein mediates virus-cell fusion, and its cleavability is directly related to virus virulence. The HN homotetramer is composed of a short N-terminal cytoplasmic followed by trans membranal, and outer membrane stalk and head domains. It recognizes sialic acid-containing receptors on the target cell surface, promotes F protein-driven virus-cell fusion activity, and acts as a neuraminidase by removing sialic acid from progeny virus particles (Huang et al. 2004).

To express the viral proteins in SJ a structure-guided analysis was performed to select epitopes vital for recognition by the immune system together with structural motifs essential to the formation of stable proteins. The designed HN protein variant 1 (HNv1) was constructed from the head fragment of native HN (Figure 4a). The designed F protein variant 1 (Fv1) was constructed from the F2 and F1 fragments connected by their natural linker (Figure 4b). A histidine tag was added to the C-terminus region of both viral subunits, for purification and detection purposes.

**Figure 4.**
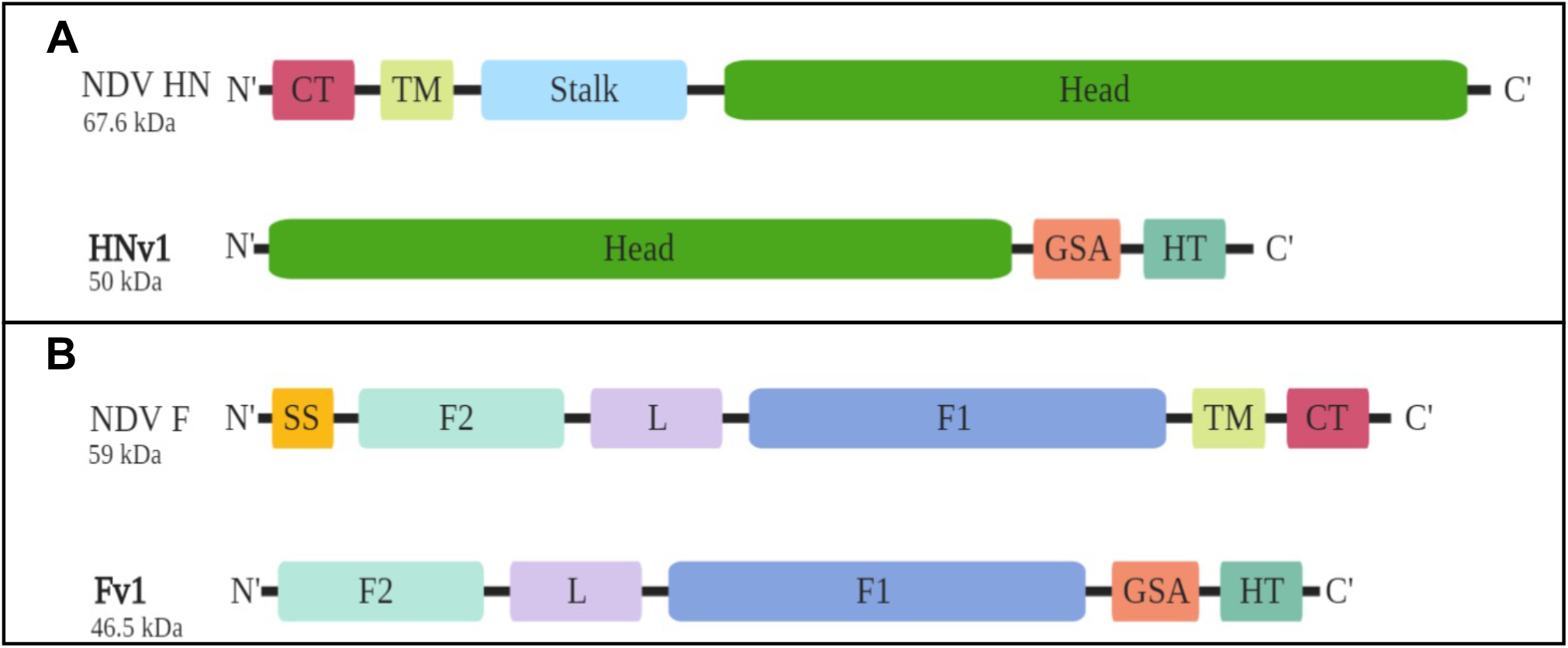
Schematic illustration of the native and designed HN and F proteins. Sequence and structural alignments were used to design NDV surface proteins suitable for expression in a prokaryotic system. **A**. Native NDV HN protein (top) and designed HNv1 polypeptide (bottom in bold). **B**. Native NDV F protein (top) and designed Fv1 polypeptide (bottom in bold). CT-ytoplasmic tail, TM-transmembrane domain, Stalk-the conjunctive segment of the tetrametric form, Head-globular fragment, GSA-link adaptor encoding for two repeats of three glycines followed by serine, HT-x6 Histidine tag, SS-signal sequence, F1/F2-segment 1 or 2 of F protein, L-linker.

The NDV protein subunits were cloned without or downstream and in-frame with SP 104. Vectors were then transformed into wild type (WT) SJ cells. The presence of SP 104 lowered the intracellular expression of Fv1 (Fv1-104) (Figure 5A). In parallel, no polypeptide secretion to the medium was detected (Figure 5B). Notably, a significant reduction in growth rate was measured after induction of Fv1-104 expression, suggesting that it may have had a toxic effect on the cell. In contrast, addition of SP 104 to HNv1 (HNv1-104) did not significantly change the intracellular expression of the target protein (Figure 5A) and induced secretion of HNv1 to the culture medium (Figure 5B). Serum collected from NDV-vaccinated chickens was able to detect HNv1 but not the Fv1 polypeptide (Figure 5C). Taken together, the expression, secretion, and immune relevance of the HNv1 polypeptide renders it a better candidate for a subunit vaccine produced in SJ, as compared to Fv1 in its present form.

**Figure 5.**
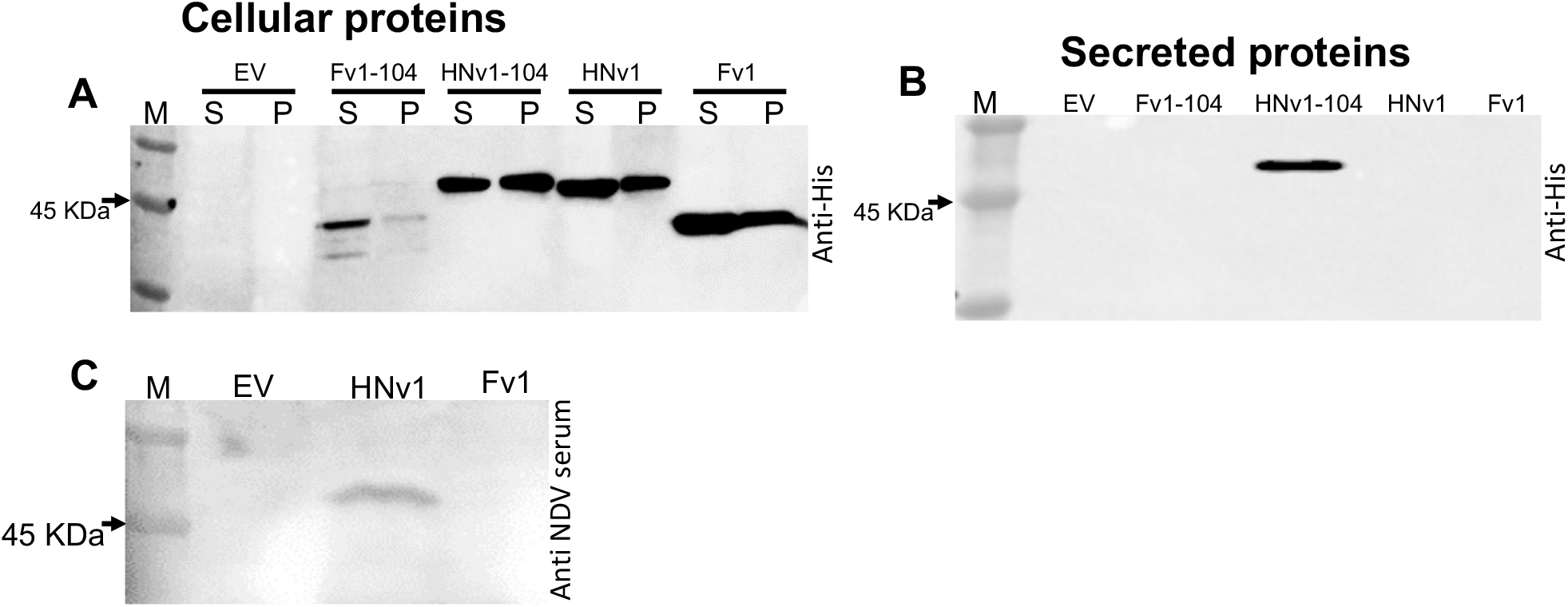
Expression, secretion and immune recognition of recombinant NDV subunits. Western blot of SJ-expressed Fv1 and HNv1 polypeptides with and without SP 104. Rabbit anti-x6 His tag-HRP or serum collected from NDV-immunized chickens and goat anti-chicken HRP were used for detection. (A) Cellular protein content after induction of both viral subunits expressed with or without SP104. S, cell lysate supernatant (soluble); P, cell lysate pellet (insoluble). (B) Detection of Fv1 and HNv1 secreted into the growth medium. (C) Proteins from the soluble cellular fraction detected with anti-NDV serum and goat anti-chicken HRP. M, protein size marker; EV, empty vector; HNv1/Fv1, the respective viral subunit; HNv1-104/Fv1-104, the respective viral subunit with the signal peptide SP 104.

### Chicken vaccination with viral subunits expressed in SJ cells

To examine the ability of HNv1 to induce a specific immune response, two doses of clarified and concentrated bacterial lysates were injected at a two-weeks interval, into three-week-old chicks. Two weeks after the second injection, sera samples from the HNv1-vaccinated group showed a small increase in NDV-specific antibodies (Figure 6) as compared to the PBS-vaccinated group, with two animals displaying antibody levels comparable to those obtained following vaccination with the commercial VH vaccine.

**Figure 6.**
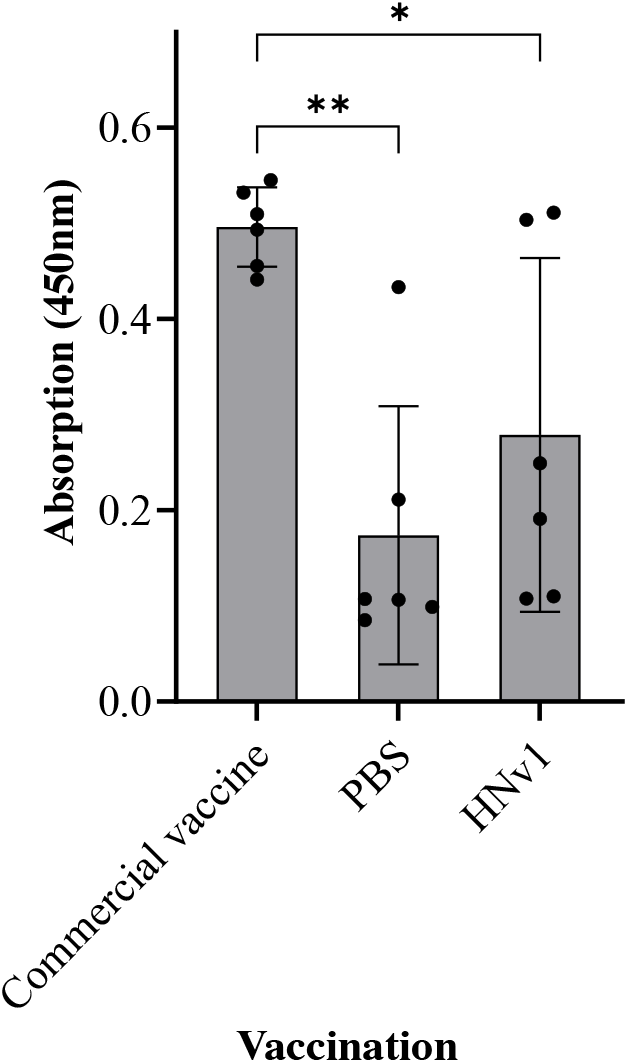
Serum antibodies level against whole NDV virus. Serum collected from chickens vaccinated with viral proteins was used in a sandwich ELISA performed as described in the material and methods. The ELISA signal is presented as mean absorption at 450 nm ± SD (n = 6) for each group. PBS was injected as negative control, Commercial vaccine of an inactivated VH strain was injected as a positive control and HNv1 as whole cell lysate of SJ expressing the polypeptide. One-way ANOVA with Tukey’s post hoc was performed, and statistically significant differences are marked with asterisks (*p<0.05, ** p<0.01).

## Discussion

The microbial world holds an enormous degree of biological diversity of which a large fraction remains untapped due to lack of appropriate molecular tools and scientific background. Unique bacterial families may carry functional properties that can facilitate various biotechnological applications limited by the characteristics of commonly applied microorganisms. *Sphingomonadaceae*, a unique LPS-free Gram-negative bacteria, belongs to *Alphaproteobacteria*, and forms a phylogenetically diverse group of environmentally abundant bacteria that have gained some attention for their potential in bioremediation and their use in the food industry (Kaczmarczyk et al. 2013). This study explored the possibility of using SJ, a member of the *Sphingomonadaceae*, as a potential platform for production of proteins, with emphasis on therapeutic applications. SJ was chosen due to its cultivability, genome sequence availability and amenability to genetic manipulation. In addition to the use of bacteria as a platform for production of therapeutic proteins, their use as a vehicle for delivery of relevant therapeutics, e.g., vaccine antigens, may also offer many advantages over traditional administration routes. Bacteria can be engineered to display or secrete target antigens and can be administered as a live vaccine to stimulate the host immune system (da Silva et al. 2014; Finn and Egan 2018; Szatraj et al. 2017). Although the live bacterial vaccine vector is a powerful adjuvant, certain disadvantages, such as safety and toxicity, must also be considered (Ding et al. 2018).

Bacteria commonly used to deliver antigens are genetically modified such that most of their pathogenic components are removed. Yet, the immunogenic LPS is toxic for embryos, restricting the usage of Gram-negative bacteria. An attractive and developing route of vaccine administration in the poultry industry is *in-ovo* vaccination, where vaccine antigens are injected to embryonated eggs prior to hatching. This method facilitates automated vaccination in the hatcheries, which reduces the need for extensive manual labor and stress to the birds (Saeed et al. 2019). Injection of live SJ *in ovo* proved safe, eliciting no signs of toxicity, in contrast to LPS-producing bacteria in which all embryos died, highlighting its potential use as a live vaccine vector. Several vaccine agents have already been shown to induce an immune response following *in ovo* administration. Nevertheless, commercial *in-ovo* vaccination is still limited to few attenuated viral vaccines (Saeed et al. 2019).

The effectiveness of subunit vaccines for a number of disease-causing viruses has been shown in humans and animals, including chickens (McAleer et al. 1984; Fingerut et al. 2003; Pitcovski et al. 2003; Pitcovski et al. 2005; Goldenberg et al. 2016). When compared to conventional vaccines, subunit vaccines bear no risk of incomplete inactivation or reversal to virulence as experienced with inactivated and attenuated vaccines, respectively. In addition, they are manufactured in safe-to-handle expression systems which can be relatively quickly adjusted to address emerging variants. Furthermore, the immune response is directed to the epitopes that confer protective immunity, whereas vaccination with intact viruses can induce high titers of non-neutralizing antibodies.

Polypeptides derived from NDV were used here as a model to evaluate the ability of SJ to express de novo designed polypeptides designed for subunit vaccination. NDV envelop proteins, including F and HN are difficult to express and produce at high quantities. Several studies had been conducted to produce these proteins in various heterologous systems, such as mammalian cell culture, plant-based systems, insect cells, yeast, virus-vector and *E. coli* (Taylor et al. 1990; Nallaiyan et al. 2010; Gu et al. 2011; Iram et al. 2014; Shahriari et al. 2015; Shahriari et al. 2016). While promising results have been reported, a cost-effective and scalable system that produces natively folded and immunogenically functional proteins is still lacking, limiting their extensive application. For a subunit viral vaccine to be functional, it is fundamental that the protein produced by the selected expression system retains its neutralizing epitopes, to induce an immune reaction to conformational epitopes which will subsequently improve virus neutralization (Khow and Suntrarachun 2012). To produce the F and HN subunits in a prokaryote-based system, codon-optimized constructs were designed based on the sequence of the proteins predicted to preserve most neutralizing epitopes for antibodies, while complex and hydrophobic segments were removed to ensure folding and prevent aggregation (Figure 4). Soluble forms of NDV-derived subunits were detected in low quantities in the cell lysate supernatant (Figure 5). In contrast, when expressed in *E. coli* BL21 (DE3), only insoluble proteins which accumulated in inclusion bodies were detected. The soluble subunit formed suggests that the SJ system facilitated correct folding of these complexed proteins. Furthermore, serum from chicken immunized with attenuated NDV, recognized SJ-expressed HNv1 (Figure 5c), suggesting that immunological epitopes were preserved. Additionally, immunization of chickens with lysate from cells of SJ expressing HNv1 induced a specific immune reaction that manifested in an overall increase in anti-viral antibody titers.

When expressed with SP 104, Fv1 was not secreted to the culture medium and a reduction in cell growth rate was noted. In contrast, addition of SP 104 to the protein subunit derived from the NH protein (HNv1-104) induced its secretion to the culture medium. These results indicate that SJ can both produce soluble and functional NDV-derived subunits and secrete them to the culture medium, albeit, in low quantities with the SP tested. Future testing of additional SPs and promoters is needed to further improve secretion and production of these polypeptides.

Despite the interest in many different physiological aspects of *Sphingomonadaceae*, tools for protein expression are still limited (Zhang et al. 2018). To expand the set of molecular tools available for protein expression in *Sphingomonadaceae*, this work developed a secretion system based on SP native to SJ. While the secretion capabilities of Gram-negative bacteria are not considered high (Burdette et al. 2018), it was theorized that *Sphingomonadaceae* have superior secretion competence compared to *E. coli* since they utilize a wide range of external compounds that require the secretion of a variety of enzymes (Byun and Blinkovsky 2013). The SecYEG complex is a conserved membrane-integrated heterotrimeric translocation channel (Saunders et al. 2006; Natale et al. 2008; Green and Mecsas 2016), responsible for most bacterial protein translocation to the periplasmic space of Gram-negative bacteria. This conserved complex can be found in several *Alphaproteobacteria* species, where it was shown to function as it does in other bacteria (Gatsos et al. 2008). SecYEG in the SJ proved 53% identical to that of the *E. coli* K 12 strain, suggesting that SJ indeed has a Sec pathway. In addition, genes encoding proteins coupled with SP potentially secreted via Sec system were identified and 6 putative SPs were tested for their ability to induce secretion of foreign proteins. The 6 SP tested imparted markedly different effects on secretion of foreign proteins by SJ. The secretion capability and efficiency of a protein carrying SP can vary dramatically, depending on the specific gene and the growth conditions. For these reasons, a secretion reporter system was designed, based on the pVH vector using two reporter genes. SP 2, SP 5, and SP 104 promoted protein secretion levels and while all three were associated with comparable αAmy secretion, SP 104 markedly increased mCherry secretion in comparison to the other SPs (Figure 3). These results highlighted that SP potency is greatly dependent on the downstream gene and on the expression conditions. Therefore, the selection of a SP sequence suited for a broad range of genes and conditions is important for development of an efficient secreted protein expression system.

To conclude, exploitation of LPS-free *Sphingomonadaceae* bacteria offers a significant advantage over conventional gram-negative bacteria expression system. This bacterial expression system successfully produced and secreted soluble complex proteins, including virus-derived vaccine polypeptides.

## Author Contribution

KE and IY conceived and designed research. KE, RBA, DE and ES conducted experiments. IB designed DNA constructs and contributed analytical tools. KE, ES, JP and IY analyzed data. KE, ES, JP and IY wrote the manuscript.

## Acknowledgement

This work was partly funded by a grant from the Chief Scientist of the Israeli Ministry of Agriculture.

## Ethics approvals and consent to participate

Animal care and ethics are in accordance with approval 020b979750 given by the ethics committee of the Kimron Veterinary Institute of Israel.

